# Oxford Nanopore MinION sequencing enables rapid whole-genome assembly of *Rickettsia typhi* in a resource-limited setting

**DOI:** 10.1101/292102

**Authors:** Ivo Elliott, Elizabeth M. Batty, Damien Ming, Matthew T. Robinson, Pruksa Nawtaisong, Mariateresa de Cesare, Paul N. Newton, Rory Bowden

## Abstract

The infrastructure challenges and costs of next-generation sequencing have been largely overcome, for many sequencing applications, by Oxford Nanopore Technologies’ portable MinION sequencer. However the question remains open whether MinION-based bacterial whole-genome sequencing (WGS) is by itself sufficient for the accurate assessment of phylogenetic and epidemiological relationships between isolates and whether such tasks can be undertaken in resource-limited settings. To investigate this question, we sequenced the genome of an isolate of *Rickettsia typhi*, an important and neglected cause of fever across much of the tropics and subtropics, for which only three genomic sequences previously existed. We prepared and sequenced libraries on a MinION in Vientiane, Lao PDR using v9.5 chemistry and in parallel we sequenced the same isolate on the Illumina platform in a genomics laboratory in the UK. The MinION sequence reads yielded a single contiguous assembly, in which the addition of Illumina data revealed 226 base-substitution and 5,856 in/del errors. The combined assembly represents the first complete genome sequence of a human *R. typhi* isolate collected in the last 50 years and differed from the genomes of existing strains collected over a 90-year time period at very few sites, and with no re-arrangements. Filtering based on the known error profile of MinION data improved the accuracy of the Nanopore-only assembly. However, the frequency of false-positive errors remained greater than true sequence divergence from recorded sequences. While Nanopore-only sequencing cannot yet recover phylogenetic signal in *R. typhi*, such an approach may be applicable for more diverse organisms.

## Introduction

Until recently, whole-genome sequencing (WGS) has been the preserve of high-income settings. Although the costs of WGS have dramatically decreased over the past decade^1^, the need for initial investment in sequencing platforms and associated equipment and for the technical expertise to run and maintain them have prevented their introduction into lower income settings. Downstream processing of sequencing data is frequently hampered by poor Internet connectivity, a lack of local expertise and the perceived requirement for substantial computational infrastructure.

Oxford Nanopore Technologies (ONT) MinION sequencer is a portable device for DNA and RNA sequencing that generates data for local analysis, in real time. The MinION weighs under 100g and plugs directly into a laptop via a USB port, with no additional computing infrastructure required^2^. Only basic laboratory facilities are needed to extract DNA and prepare sequencing libraries. The MinION has been used to sequence viral^3^, bacterial^4,5^ and eukaryotic^6–9^ genomes. In spite of its high raw error rate, Nanopore data can in many cases produce highly contiguous assemblies. Illumina short-read data can be added to improve consensus accuracy, in many cases to that of a high-quality draft assembly, without extra finishing steps.

The Lao People’s Democratic Republic (Laos) is a landlinked country of ~7 million people in Southeast Asia. Laos is ranked 135^th^ in the world on the Human Development Index (HDI), a composite statistic of life expectancy, education and per capita income^10^. There are few functioning molecular biology laboratories in Laos and no previous published WGS projects.

*Rickettsia typhi* is an obligate intracellular, Gram-negative bacterium in the family Rickettsiaceae that causes the disease murine typhus and has a worldwide tropical and sub-tropical distribution. The pathogen is transmitted to humans primarily by flea faecal contamination of the bite of the Oriental rat flea *Xenopsylla cheopis.* The disease is an important and grossly under-recognized global cause of febrile illness. Those affected suffer from high fever, headache, myalgia, arthralgia, nausea and vomiting and may have a macular rash. Complications are infrequent but include myocarditis, meningoencephalitis, seizures and renal failure^11^. When diagnosed, recovery is typically rapid after treatment with doxycycline and mortality is low (around 1-2%) with antibiotic treatment^12^.

Despite its importance as a pathogen, to date just three *R. typhi* whole-genome sequences, with wide geographic and temporal distribution, have been published. The type-strain, Wilmington, was isolated from a patient in North Carolina in 1928 and the sequence was published in 2004^13,14^. Two further sequences, originating from a patient in Northern Thailand collected in 1965 and the other from a bandicoot rat, *Bandicota* sp., collected in Burma (now Myanmar) in the 1970s ^15–17^ were published online in 2014 (Table 1).

**Table 1.** Strain information. The Wilmington, B991CWPP, and TH1527 strains are previously assembled strains of R. typhi; TM2540 is the assembly produced in this study.

| Strain | Source | Accession | Genome length (bp) | Predicted genes | GC percentage |
| --- | --- | --- | --- | --- | --- |
| Wilmington | Human, North Carolina, USA. 1928. | NC_006142.1 | 1111496 | 865 | 0.289 |
| B9991CWPP | Bandicoot rat, Myanmar. 1970s | NC_017062.1 | 1112957 | 865 | 0.289 |
| TH1527 | Human, Chiang Rai, Thailand. 1965 | NC_017066.1 | 1112372 | 865 | 0.289 |
| TM2540 | Human, Bolikhamxay, Laos. 2012 | ERZ497871 | 1111939 | 866 | 0.289 |

We report the successful whole genome sequencing of a human isolate of *R. typhi* using only the MinION platform in Laos. We assess the potential for use of ONT data alone to perform comparative analyses, and discuss the challenges of undertaking WGS in resource-limited settings.

## Materials and Methods

### *R. typhi* culture and DNA extraction

Frozen L929 mouse fibroblast cells (ATCC CCL-1), infected with *R. typhi* isolated from the blood of a patient, TM2540, from Pakkading District (~18.33° N 104.0° E), Bolikhamxay Province, Laos with suspected acute murine typhus infection presenting in 2012 to Mahosot Hospital, Vientiane, were reanimated. Frozen aliquots were briefly thawed at room temperature and then transferred to 25 cm^2^ cell culture flasks containing a L929 cell monolayer at 80% confluence in RPMI 1640 medium (Gibco, USA), supplemented with 10% foetal calf serum (Sigma-Aldrich, USA). Flasks were incubated at 35 °C in 5% CO_2_ atmosphere for seven days, then cells were mechanically detached and transferred to a 75 cm^2^ flask and again incubated at 35 °C in 5% CO_2_ atmosphere.

For DNA extraction, cells from three 75 cm^2^ flasks were mechanically detached, re-suspended and transferred to a 50 ml conical-bottom centrifuge tube and centrifuged at 3,220*×g* for 10 minutes. The pellet was re-suspended in 3ml fresh medium and transferred to 1.5 ml micro-centrifuge tubes. Tubes were vortexed for one minute, then centrifuged at 300*×g* for 3 minutes. The supernatant was passed through a 2 μm filter (Corning, USA), then mixed with 10 μl/ml DNAase (1.4 μg/μl) and incubated at room temperature for 30 minutes. The mixture was centrifuged at 18,188×*g* for 10 minutes and washed twice with 0.3 M sucrose (Sigma-Aldrich, USA). DNA was extracted from the *R. typhi* pellet using the DNeasy Blood & Tissue kit (Qiagen, USA) and eluted products were stored immediately at −20 °C.

DNA was quantified using Qubit dsDNA High Sensitivity assay kit (Thermofisher, USA) following manufacturer’s protocols and assayed for *R. typhi* sequences by quantitative polymerase chain reaction (qPCR) targeting the 17kDa outer membrane antigen^18^.

### Preparation of MinION sequencing libraries

MinION sequencing libraries were produced using the ONT 1D Genomic DNA by ligation (SQK-LSK108) protocol. Briefly, 1.4 μg of DNA was fragmented in a Covaris g-TUBE (Covaris Ltd., Brighton, U.K.) by centrifugation. Sheared DNA was repaired using NEBNext FFPE repair mix (New England Biolabs, MA, USA). End-repair and dA-tailing were performed with the NEBNext Ultra II End Repair/dA-tailing module. Adapter ligation used the NEB Blunt/TA Ligase Master Mix, and the library was purified using Agencourt AMPure XP beads (Beckman Coulter Inc., CA, USA).

### MinION sequencing in Laos

MinION libraries were sequenced for 48 hours on an ONT MinION R9.5 flow cell, connected to a Dell Latitude E5470 XCTO laptop with 256GB SATA Class 20 solid state drive. ONT fast5 data files were base-called using the ONT Albacore module.

### Illumina sequencing in Oxford

An Illumina sequencing library was generated from the same sample of *R. typhi* DNA using the Nextera (Illumina) library preparation method. The library was sequenced on the Illumina MiSeq with 2×260bp reads. A total of 1,201,068 read pairs were sequenced, giving 312Mbp of total sequence data.

### Bioinformatic analysis

The species composition of the MinION reads was identified using Centrifuge software^19^ testing against the prebuilt non-redundant database and keeping only the best hit per read.

Assembly strategies that used MinION data alone, or in combination with Illumina short reads, were assessed. A draft genome assembly was generated from the MinION reads using Canu^20^ with the suggested parameters for ONT sequencing “correctedErrorRate=0.120-nanopore-raw” and an estimated genome size of 1.1Mb, and polished using Nanopolish^4^. The Illumina reads were mapped to the Canu+Nanopolish assembly using bwa 0.7.12^21^ and the mapped reads were used to error-correct the assembly using Pilon vl.22^22^. To check for enrichment of errors in regions of low coverage, the ONT reads were mapped onto the corrected assembly using bwa mem to determine coverage and the coverage at error positions was compared to the total coverage distribution. To look for enrichment of errors near homopolymers, we compared the distance to the nearest homopolymer of 5bp or longer in length for each corrected error to a random sample of positions in the genome. BUSCO v3 using the OrthoDB version 9 bacterial dataset was used to assess gene content^23^. The Illumina short reads alone were assembled using Unicycler^24^. Prokka vl.14^25^ was used to annotate both the new assemblies and the existing *R. typhi* genomes to give consistent data for comparison. The Wilmington reference strain was used to train the gene model for the Prodigal gene prediction used in Prokka, and this training set was used to annotate the other three genomes. The new genome was rotated to begin with the *yqfL* gene for consistency with other *R. typhi* genomes.

### Data availability

The three existing *R. typhi* genomes were downloaded from RefSeq under the accession numbers NC_006142.1, NC_017066.1 and NC_017062.1. The sequence data is available at the ENA under project PRJEB2567. The new strain is named TM2540 and the assembly is available at the ENA under accession ERZ497871.

## Results

MinION flow cells and reagents were shipped from Oxford, United Kingdom to Laos at +4°C. On dispatch 1,242 active pores were available and on receipt after 72hrs travel, 1,103 active pores were available. Sequencing using the MinKNOW platform was performed for 48 hours and generated approximately 250,000 reads, with a total fast5 file size of 20GB. 222,848 reads passed quality filters and were used in further analysis.

We assessed the species composition of the reads which passed quality filters. Since the *Rickettsia typhi* sample was grown in L929 fibroblast cells, we expected to see some contaminating reads from the mouse genome. 134,802 (60%) of the reads were classified as belonging to the genus *Rickettsia.* 126,447 reads were classified as *Rickettsia typhi* at the species level and 41,428 (19%) were classified as belonging to the genus *Mus* (Figure 1).

**Figure 1.**
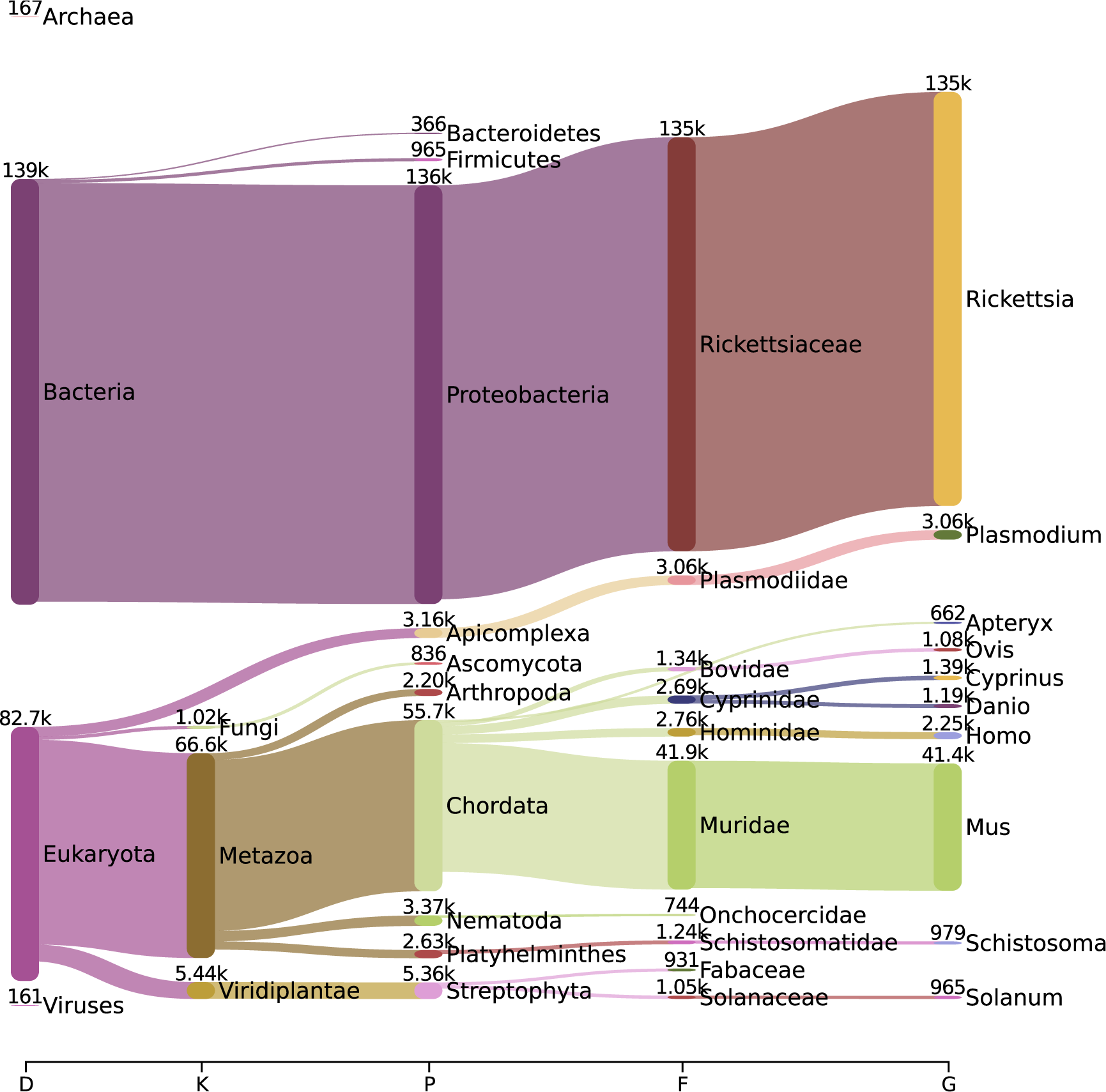
Pavian visualization showing the proportions of ONT reads assigned up to the genus level^42^.

The FASTQ files produced by Albacore were assembled using Canu, an assembler designed for noisy single-molecule sequencing such as that produced by the MinION. This produced a single-contig assembly of l,078,916bp. Nanopolish, a tool to use the signal-level data from Oxford Nanopore sequencing, was used to improve the consensus sequence of the Canu assembly, producing a new assembly of l,099,322bp. To further correct errors in the assembly produced from long reads alone, we polished the Canu+Nanopolish genome assembly using Pilon, which uses short read sequencing data to correct errors. We repeated rounds of mapping and correction with Pilon until no further errors were corrected. After 4 rounds of correction, Pilon did not correct any further errors, with the exception of two short indels which were removed and then reinserted by successive rounds of polishing. 94.2% of errors were corrected by the first round of polishing, with a further 4.8% corrected in the second round (Supplementary Table 1).

After this process, a total of 6,082 errors were corrected, with over 95% (5,856) being small insertions and deletions, with the result that 12kb was added to the genome during polishing. In comparison, when Pilon was run on the Canu draft genome before Nanopolish polishing, 19,970 errors were corrected, confirming that Nanopolish improves the draft genome.

We assessed the ONT read coverage and proximity to a homopolymer run (5 bases or longer) for each of the errors we corrected with Pilon. We observed no difference in the coverage distribution at sites with errors compared to the overall coverage distribution (Supplementary Figure 1). However, the positions around homopolymer runs were enriched for errors (Figure 2) - 15% of the final polished genome is within 5bp of a homopolymer run (165kb), while 43% of errors (2,641) fall into these regions.

**Figure 2.**
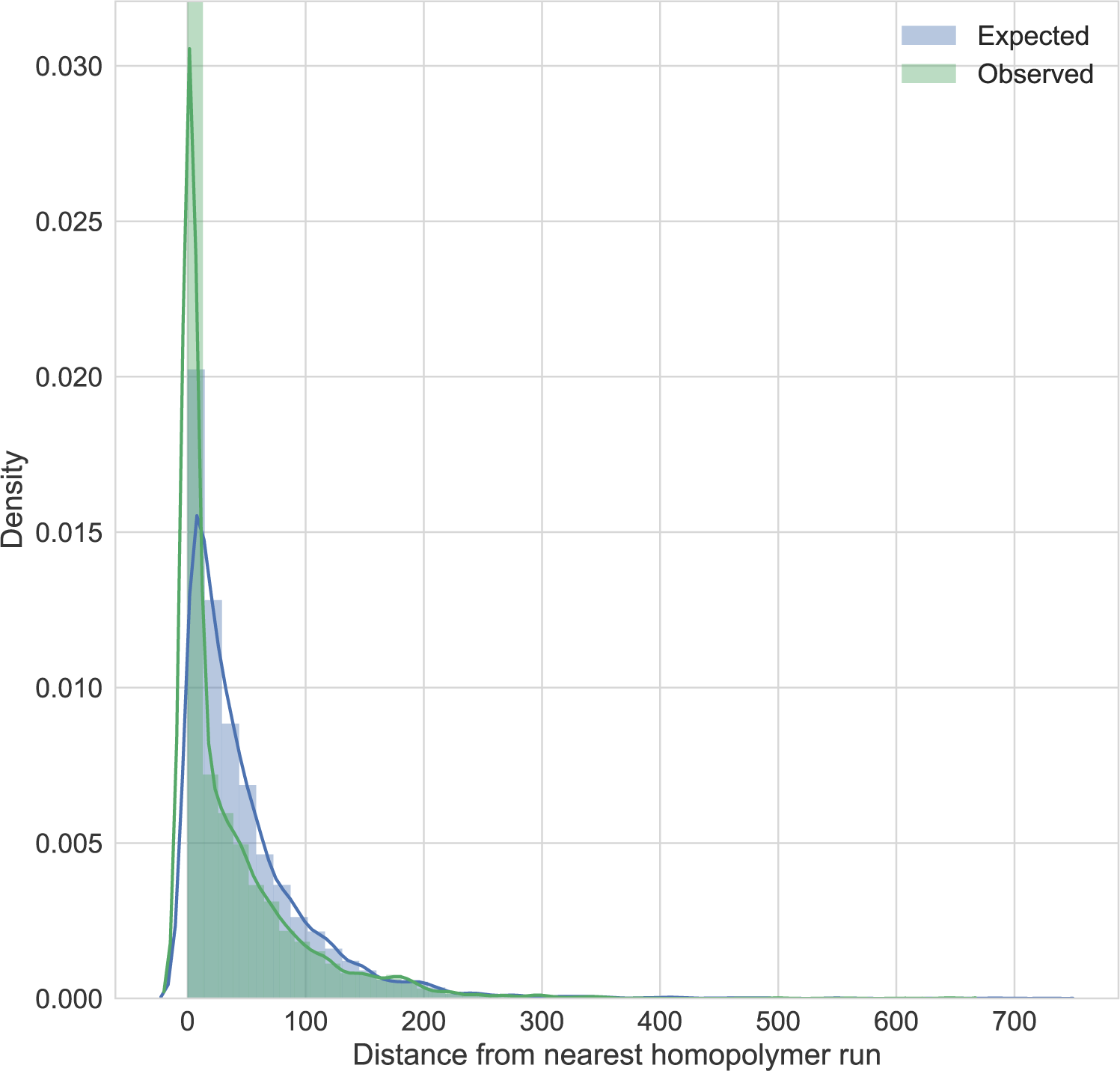
The observed distance from a homopolymer run for the errors corrected by Pilon in the draft genome, compared to the expected distribution if errors were randomly distributed across the genome.

The Illumina data alone was assembled using Unicycler to compare with assemblies using ONT only and the combined ONT and Illumina data. The assembly generated by Unicycler had 29 contigs with a total length of 1,161,002bp, and an n50 of 166kb. However, this included 20 short contigs under 6kb, which are closely related to the mouse genome by BLAST search. These contigs are identified by their low coverage depth relative to *Rickettsia typhi* contigs. After removal of contaminating mouse contigs, the Unicycler assembly has 8 contigs with a total length of 1,111,455bp.

BUSCO was used to quantitatively measure the genome completeness of our assemblies based on a set of 148 conserved single-copy genes (Table 2). None of the genes were found in the initial Canu assembly. Inspection of the TBLASTN results shows that while there were hits to many of the conserved genes, they fall below the threshold where BUSCO will call them as present. After polishing with Nanopolish using ONT data only, 14.2% of the genes could be found as a complete copy, with a further 32.4% found as fragmented genes. Two rounds of Pilon polishing improves this to 88.5% of genes found as complete copies. Running BUSCO on the Wilmington reference strain gives the same number of complete genes, suggesting this is the complete set of the orthologous reference genes that exists in *R. typhi.* The Unicycler assembly also contains this complete set of genes.

**Table 2.**
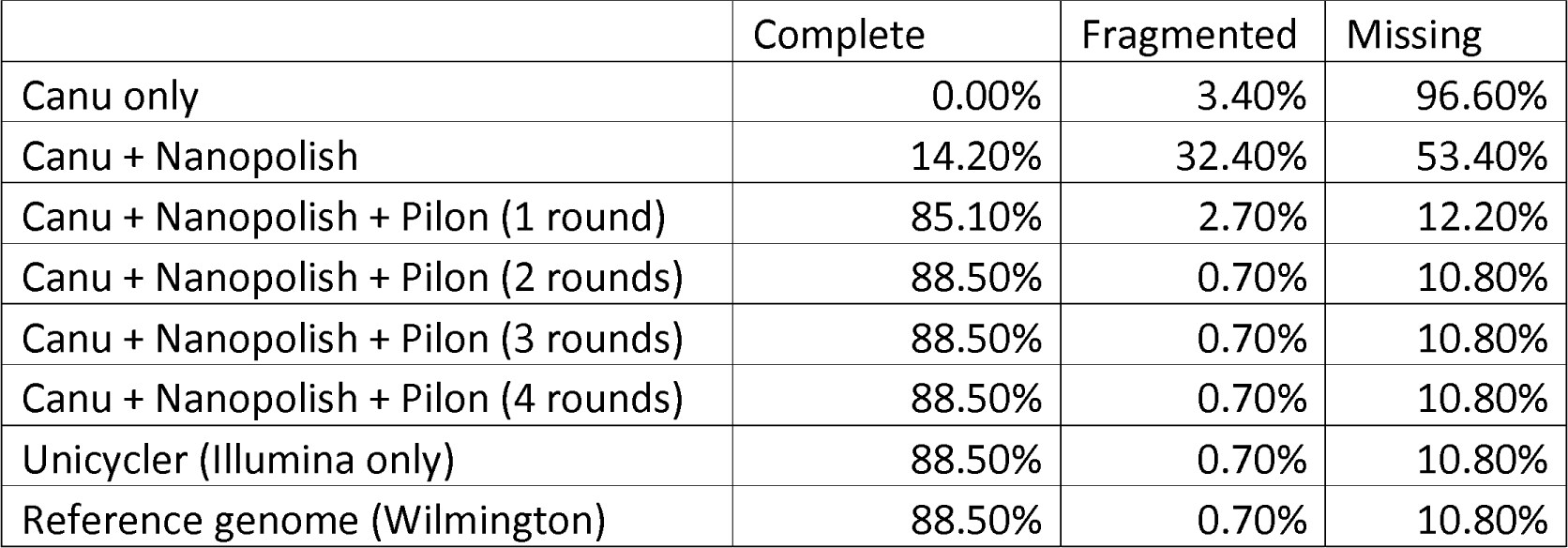
Proportion of BUSCO bacterial gene set found as complete genes, fragmented genes, or missing for different assemblies of the TM2540 strain, and the reference Wilmington assembly. All complete genes were single-copy.

The final polished genome assembly is 1,111,939bp, extremely similar to the three available genomes, which range from 1,111,496-1,112,957bp in length (Table 1). Alignment of the four whole genomes in Mauve^26^ shows that the genomes are co-linear and that no rearrangements have taken place between strains (Supplementary Figure 1).

The TM2540 genome contained 866 predicted genes, very close to the 865 genes predicted in all other strains. Roary was used to define the core genome (genes present in all strains at 90% identity) of the four *R. typhi* strains, which included 863 genes. Of the 5 accessory genes not found in all strains, two were found in three of the four strains, and three were unique to a single strain. Two of the genes not found in all samples were the same gene, but clustered separately due to a 150bp insertion in the B991CWPP strain that caused the sequence to fall below the 90% identity threshold to cluster with the other three strains. In two of the five accessory genes, a single base insertion in one strain has created a premature stop codon, splitting a gene into two sequences in that strain. In the final case, the gene is present in three strains next to a duplicate pseudogene which has lost the start and end of the gene, while in the fourth strain a deletion has removed the end of the pseudogene and the start of the gene, leaving two incomplete and presumably inactive copies of the gene. All of the 5 variably present genes are annotated as hypothetical genes.

Table 3 shows the SNP differences between strains determined from a whole genome alignment, with 15-23 SNPs separating each strain.

**Table 3.** Pairwise differences between strains based on the whole genome alignment.

|  | Wilmington | B9991CWPP | TH1527 | TM2540 |
| --- | --- | --- | --- | --- |
| Wilmington | 0 | 23 | 17 | 22 |
| B9991CWPP | 23 | 0 | 16 | 21 |
| TH1527 | 17 | 16 | 0 | 15 |
| TM2540 | 22 | 21 | 15 | 0 |

We compared the results of the final polished assembly to the best assemblies generated using Illumina data alone (filtered Unicycler assembly) or ONT data alone (Canu + Nanopolish assembly). The Illumina assembly has 866 predicted genes, and the core genome using this assembly has 862 core genes and 7 accessory genes. This includes one gene unique to the Illumina assembly, which is part of a gene split into two pieces at a contig boundary. The ONT assembly has 1745 predicted genes, with many genes duplicated in tandem, likely due to sequence errors, and the core genome using this assembly has only 774 genes.

## Discussion

WGS of pathogens is an increasingly important tool in both research and clinical settings, whose use has increased over the last decade as its cost has decreased and availability improved. However, the initial investment in sequencing and computing equipment and the technical expertise needed to produce and analyse data remain a barrier to its introduction into resource-poor settings. Laos has very few clinical or research laboratories and relatively poor Internet connectivity^27^. Although our research laboratory can perform molecular diagnostics and cell culture, to date there has been no sequencing capacity, necessitating the slow and costly shipment of samples to other countries to undertake sequencing projects. The portable nature of the MinION sequencer has allowed the first bacterial WGS to be performed in Laos. We were able to run all steps of our bioinformatic analysis (except Centrifuge) on a laptop computer (MacBook Pro 2017) without continuous Internet access, demonstrating that sequencing and analysis can take place in relatively remote settings.

Murine typhus is an important and severely neglected tropical and subtropical disease of worldwide distribution. In Vientiane an estimated 10% of non-malarial fevers in adult inpatients is caused by *R. typhi*^28^. Surprisingly, for a disease to which many millions of people are potentially exposed, there are only three published whole genome sequences, of which the last human isolate dates back more than 50 years.

The *R. typhi* genome is 1.1Mb and contains almost no repeats, which makes genome assembly for this species relatively simple. Running Canu with the default ONT parameters was sufficient to assemble the genome into a single contig and we polished the assemblies using both Nanopolish, to utilize the signal-level data from ONT sequencing, and Pilon, to use the extra information available from Illumina sequencing with its lower error rate. While its synteny with other genomes suggests that there are no large misassemblies in our genome, the two methods of polishing corrected many small SNP and indel errors in the ONT only draft genome. Thus even for relatively simple bacterial genomes, a combination of ONT with data from another technology with a low error rate is currently necessary to produce an accurate sequence. In the case of *R. typhi*, the strains are so closely related that to distinguish between them requires a highly accurate sequence, and using ONT data alone the sequencing errors are more numerous than the true differences between strains. Removing regions of the genome close to homopolymer runs would remove a large number of positions with errors by filtering out a comparatively small amount of the genome, but would still leave over 2,000 more errors than true differences. ONT data alone may suffice for applications which can work with sequences which still retain some base-pair level errors, such as determining large-scale genome rearrangements^29^, determining the species of an unknown sample^30,31^, or detecting antimicrobial resistance genes^32^. Increases in sequencing accuracy, combined with bioinformatics advances in assembly and polishing software, may allow for the future use of ONT data alone to give complete and accurate sequences, increasing the applications for this device.

Comparative analysis of the four *R. typhi* genomes shows very little variation between strains. Very few genes are variably present between strains, and gene order is completely conserved. Although these four samples were collected from humans and a rodent over a long timeframe, we see only 23 SNPs between the two most divergent strains, no large-scale genome rearrangements, and very few indels. The low number of differences suggests that whole-genome sequencing may not be a useful tool to analyse the relationships between strains of *R. typhi,* as there may be few differences between strains and no recent shared ancestry.

Some limitations remain on the use of ONT’s MinION platform. Currently individual flow cells remain costly at USD500-USD900 each^33^, depending on the number purchased. However, the cost per sample can be reduced since multiplex sequencing now allows for multiple bacterial strains to be sequenced on the same flow cell^34^. Ongoing improvements in the technology continue to increase overall output and per-run capacity. Novel technologies are likely to improve access to sequencing when computational resources and support are limited and allow more cost-efficient sequencing of single samples^35^.

MinION instruments continue to be used in an ever-widening set of challenging circumstances^3,36–41^. Here we demonstrate the current applicability of the MinION to resource-poor settings where some laboratory infrastructure exists, but WGS capacity is unavailable. With concerns in some countries about the export of biological samples for WGS in other countries, MinION systems could facilitate countries without current WGS facilities to undertake such work.

## Acknowledgments

We are very grateful to Associate Professor Bounthaphany Bounxouei, Director of Mahosot Hospital, the staff of the Microbiology Laboratory, LOMWRU and wards, Assistant Professor Chanphomma Vongsamphan, Director of Department of Health Care, Ministry of Health, and H.E. Professor Bounkong Syhavong, Minister of Health, Laos, for their help and support; Narongchai Tongyoo and Phonepasith Panyanouvong for growing the isolate in cell culture; Duncan Parkes for assistance with performing base-calling.

## Author contributions

